# Analysis of the role of Nidogen/entactin in basement membrane assembly and morphogenesis in *Drosophila*

**DOI:** 10.1101/346270

**Authors:** Jianli Dai, Beatriz Estrada, Sofie Jacobs, Besaiz J. Sánchez-Sánchez, Jia Tang, Mengqi Ma, Patricia Magadan, José C. Pastor-Pareja, María D. Martín-Bermudo

**Affiliations:** School of Life Sciences, Tsinghua University, Beijing 100084, China; CABD (CSIC-Universidad Pablo de Olavide-JA), Seville 41013, Spain; Rega Institute, KU Leuven, Leuven 3000, Belgium; Randall Centre for Cell and Molecular Biophysics, King’s College London, London SE5 9AP, UK; School of Forestry and Biotechnology, Zhejiang Agriculture and Forestry Univesity, Hangzhou 311300, China

## Abstract

Basement membranes (BMs) are thin sheet-like specialized extracellular matrices found at the basal surface of epithelia and endothelial tissues. They have been conserved across evolution and are required for proper tissue growth, organization, differentiation and maintenance. The major constituents of BMs are two independent networks of Laminin and Type IV Collagen interlinked by the proteoglycan Perlecan and the glycoprotein Nidogen/entactin (Ndg). The ability of Ndg to bind in vitro Collagen IV and Laminin, both with key functions during embryogenesis, anticipated an essential role for Ndg on morphogenesis linking the Laminin and Collagen IV networks. This was supported by results from in vitro and cultured embryonic tissues experiments. However, the fact that elimination of Ndg in *C. elegans* and mice did not affect survival, strongly questioned this proposed linking role. Here, we have isolated mutations in the only Ndg gene present in *Drosophila*. We find that while, similar to *C.elegans* and mice, *Ndg* is not essential for overall organogenesis or viability, it is required for appropriate fertility. We also find, alike in mice, tissue-specific requirements of *Ndg* for proper assembly and maintenance of certain BMs, namely those of the adipose tissue and flight muscles. In addition, we have performed a thorough functional analysis of the different Ndg domains in vivo. Our results support an essential requirement of the G3 domain for Ndg function and unravel a new key role for the Rod domain in regulating Ndg incorporation into BMs. Furthermore, uncoupling of the Laminin and Collagen IV networks is clearly observed in the larval adipose tissue in the absence of Ndg, indeed supporting a linking role. In light of our findings, we propose that BM assembly and/or maintenance is tissue-specific, which could explain the diverse requirements of a ubiquitous conserved BM component like Nidogen.

**Author Summary:** Basement membranes (BMs) are thin layers of specialized extracellular matrices present in every tissue of the human body. Its main constituents are two networks of Laminin and Type IV Collagen linked by Nidogen (Ndg) and proteoglycans. They form an organized scaffold that regulates organ morphogenesis and function. Mutations affecting BM components are associated with organ dysfunction and several congenital diseases. Thus, a better comprehension of BM assembly and maintenance will not only help to learn more about organogenesis but also to a better understanding and, hopefully, treatment of these diseases. Here, we have used *Drosophila* to analyse the role of Ndg in BM formation *in vivo*. Elimination of Ndg in worms and mice does not affect survival, strongly questioning its proposed linking role, derived from *in vitro* experiments. Here, we show that in the fly Ndg is dispensable for BM assembly and preservation in many tissues, but absolutely required in others. Furthermore, our functional study of the different Ndg domains challenges the significance of some interactions between BM components derived from in vitro experiments, while confirming others, and reveals a new key requirement for the Rod domain in Ndg function and incorporation into BMs.

## Introduction

Basement membranes (BM) are specialized thin extracellular matrices underlying all epithelia and endothelia and surrounding many mesenchyme cells. This thin layer structure, which appears early in development, play key roles in the morphogenesis, function, compartmentalization and maintenance of many tissues (1).

All BMs contain at least one member of the laminin, Type IV collagen (Col IV), proteoglycan agrin and perlecan and nidogen (entactin) families. Nidogen is a 150-KDa glycoprotein highly conserved in mammals, *Drosophila*, *Caenorhabditis elegans* (*C. elegans*) and ascidians (2) (3). Nidogens have been proposed to play a key role in BM assembly by providing a link between the laminin and Col IV networks and by integrating other ECM proteins, such as perlecan, into this specialized extracellular matrix (4-7). While invertebrates possess only one nidogen, two nidogen isoforms, Nid1 and Nid2, have been identified in vertebrates. The individual knock out of either *Nid1* or *Nid2* in mice does not affect BM formation or organ development (8-10). In fact, these *Nid1* or *Nid2* null animals appear healthy, fertile and have a normal life span. However, simultaneous elimination of both isoforms result in perinatal lethality, with defects in lung, heart and limb development, which are not compatible with postnatal survival (11) (12). In addition, BMs defects are only observed in certain organs, which strongly suggest tissue-specific roles for nidogens in BM assembly and function (11). Like in mice, loss of the only nidogen gene in *C. elegans*, NID-1, is viable with minor defects in egg laying, neuromuscular junctions and position of longitudinal nerves, but not defects in BM assembly (13) (14) (15). All together, these studies reveal that Nidogens may play important roles in specific contexts, consistent with their evolutionary conservation. However, the different requirements for nidogens in BM assembly and organogenesis in mice and *C. elegans* suggest that new functions may have arisen in vertebrates. The isolation of mutants in nidogen in other organisms will help to shed light on the role of the Nidogen proteins in vivo and its conservation throughout evolution.

All nidogens comprise three globular domains, namely G1, G2 and G3, one flexible linker connecting G1 and G2 and one rod-shaped segment separating G2 and G3 domains composed primarily of endothelial growth factor repeats (16) (4, 17). A vast variety of studies using recombinant fragments of Nidogens and several binding and X-ray assays have provided a wealth of information on the binding properties of the different Nidogen domains in vitro. Thus, the globular domain G3 mediates connections to Laminin networks (4, 7, 17-20). Furthermore, the Nidogen-binding site of Laminins has accurately been mapped to a single laminin-type epidermal growth factor-like (LE) module of the laminin γ1 chain. Binding epitopes to collagen IV and Perlecan have been mapped to domain G2 (4, 5, 7, 17, 21). However, besides this substantial amount of information on the binding capacities of the different domains, the relevance of these interactions in vivo remains to be fully established. Some functional studies have been performed in cell or organ cultures. Thus, antibodies against the LE module delay BM formation in organ cultures of embryonic kidney, salivary glands and lungs (22) (23. Nevertheless, some of the predictions made from the results obtained from cell culture and in vitro systems do not always hold when tested in model organisms. For example, specific ablation in mice of the nidogen-binding site of laminins results in kidney and lung developmental defects {Willem, 2002 #1158). In contrast, deletion of the whole G2 domain in *C.elegans* is viable and does not affect organogenesis (13). These discrepancies clearly challenge a role for Nidogen as a linker between the Collagen IV and Laminin networks and emphasize the need for the use of more and diverse animal models to dissect the role of nidogens in vivo.

*Drosophila* BMs are analogous to the vertebrate ones (24). They cover the basal surface of all epithelia and endothelia and surround most organs and tissues, including muscles and peripheral nerves. Even though the composition might vary according to tissues and developmental stages, all *Drosophila* BMs contain Col IV, Laminins, Perlecan and Nidogen. However, in contrast to the 3 Col IV, 16 laminins, and 2 nidogens found in humans, *Drosophila* only produces 1 Col IV, 2 distinct Laminins and 1 Nidogen (Ndg). The reduced number of ECM components, which limits the redundancy between them, and their high degree of conservation with their mammalian counterparts, makes *Drosophila* a perfect model system to dissect their function in vivo. *Drosophila* Col IV has been identified as a homolog of Type IV Collagen in mammals, which is a long helical heterotrimer that consists of two α1 chains and one α2 chain, encoded by the genes *Collagen at 25 C* (*Cg25C*) and *Viking* (*Vkg*), respectively (25-27). The C terminal globular non-collagenous (NC1) domain and the N terminal 7S domain serve to form Col IV network (28). Loss of function mutations in either of the two *Col IV* genes in flies affects muscle development, nerve cord condensation, germ band retraction and dorsal closure, causing embryonic lethality (29). In addition, mutations in Col IV have also been associated with immune system activation, intestinal dysfunction and shortened lifespan in the *Drosophila* adult (30). Finally, while Col IV deposition in wing imaginal discs and embryonic ventral nerve cord BMs is not required for localization of Laminins and Nidogens, it is essential for Perlecan incorporation (31, 32). The *Drosophila* Laminin αβγ trimer family consists of two members comprised of two different α subunits encoded by *LamininA* and *wing blister*, one β and one γ subunit encoded by *LamininB1* and *LamininB2*, respectively (33). Same as Col IV, Laminins can also self-assemble into a scaffold through N-terminal (LN) domains interaction in short arms (34). While, elimination of laminins in *Drosophila*, in contrast to mice, does not impair gastrulation, it affects the normal morphogenesis of most organs and tissues, including the gut, muscles, trachea and nervous system (35) (36). In addition, abnormal accumulation of Col IV and Perlecan was observed in laminins mutant tissues (35). Perlecan, encoded by the *trol* (*terribly reduced optic lobes*) gene, is subdivided into five distinct domains. Interactions with Laminins and Col IV occur through domains I and V (reviewed in (37). Mutations in *trol* affect postembryonic proliferation of the central nervous system, plasmatocytes and blood progenitors (38-40). Loss of *trol* also affects the ultrastructure and deposition of laminins and Col IV in the ECM around the lymph gland (40). All together, these results suggest that the BMs components laminin, Col IV and Perlecan are all essential for proper development. In addition, they also reveal that there is a hierarchy for their incorporation into BMs that seems to be tissue-specific and required for proper BM assembly and function. However, the role of *Ndg* in *Drosophila* morphogenesis and BM assembly has remained elusive. This may be in part due to the lack of mutations in this gene.

In this work, we have dissected the role of *Ndg* in *Drosophila*. Using a newly generated anti-Ndg antibody, we have shown Ndg accumulates in the BMs of embryonic, larval and adult tissues. By isolating several mutations in the single *Drosophila Ndg* gene, we find that while, similar to *C. elegans* and mice, *Ndg* is not required for overall organogenesis or viability, it is required for fertility. Also similar to the tissue-specific defects in mice and *C.* elegans, we find that the BM surrounding the larval fat body and flight muscles of the notum is disrupted in the absence of *Ndg*. Furthermore, we observed uncoupling of Laminin and Collagen IV in the fat body of *Ndg* mutants, indeed supporting a role of Ndg as a linker between the two networks. In addition, we have performed a thorough functional analysis of the different Ndg domains in vivo, which, on one hand, supports an essential requirement of the G3 domain for Ndg function and, on the other hand, uncovers a new key role for the Rod domain in regulating Ndg incorporation into BMs. Finally, we find that BM assembly is not universal but differs depending on the tissue and propose that this could explain the diverse requirements of a ubiquitous conserved BM component like Nidogen.

## Results

### Nidogen localizes to the BM of embryonic, larval and adult tissues

Previous analysis has shown that, during embryogenesis, Ndg is expressed in multiple mesodermal cell types, such as visceral mesoderm, somatic muscle founder cells, a subset of pericardial and cardial cells and at the edges of the visceral mesoderm (41) (42, 43). Here, we decided to further analyse Ndg expression in embryonic, larval and adult tissues. In order to do this, an antibody against a peptide encoded by exon 7 was developed (see Materials and Methods). We found that in addition to the pattern described previously, similar to laminins (35), Ndg was also detected in the BM surrounding most embryonic tissues, including muscles, gut and ventral nerve cord (Fig 1A, A’, B, B’), in stage 16 embryos. However, in contrast to laminins, Ndg was not enriched at muscle attachment sites (Fig 1A). In addition, a careful analysis of Ndg expression in stage 13 embryos revealed a dotted pattern along the visceral mesoderm, which differs from the continuous line observed around the muscles or the ventral nerve cord (Fig 1C, C’). At this stage, caudal visceral mesodermal cells migrate over the visceral mesoderm. In fact, using a marker for these cells, croc-lacZ (44), we found that Ndg accumulated around them as they migrate (Fig 1C, C’). In this case, Ndg seem to be organized in track-like arrays, similar to the distribution of laminins around migrating hemocytes. Ndg was also found in migrating hemocytes, as visualized using a version of Nidogen tagged with superfolder GFP (sGFP), expressed from a duplication of the *Ndg* genomic region (Fig 1D, (45). Finally, Ndg was also found at high levels in chordotonal organs (Fig 1A, A’, asterisk). These results suggest that as it is the case for laminins, Ndg can be deposited and/or assembled in different patterns throughout embryogenesis.

**Fig 1.**
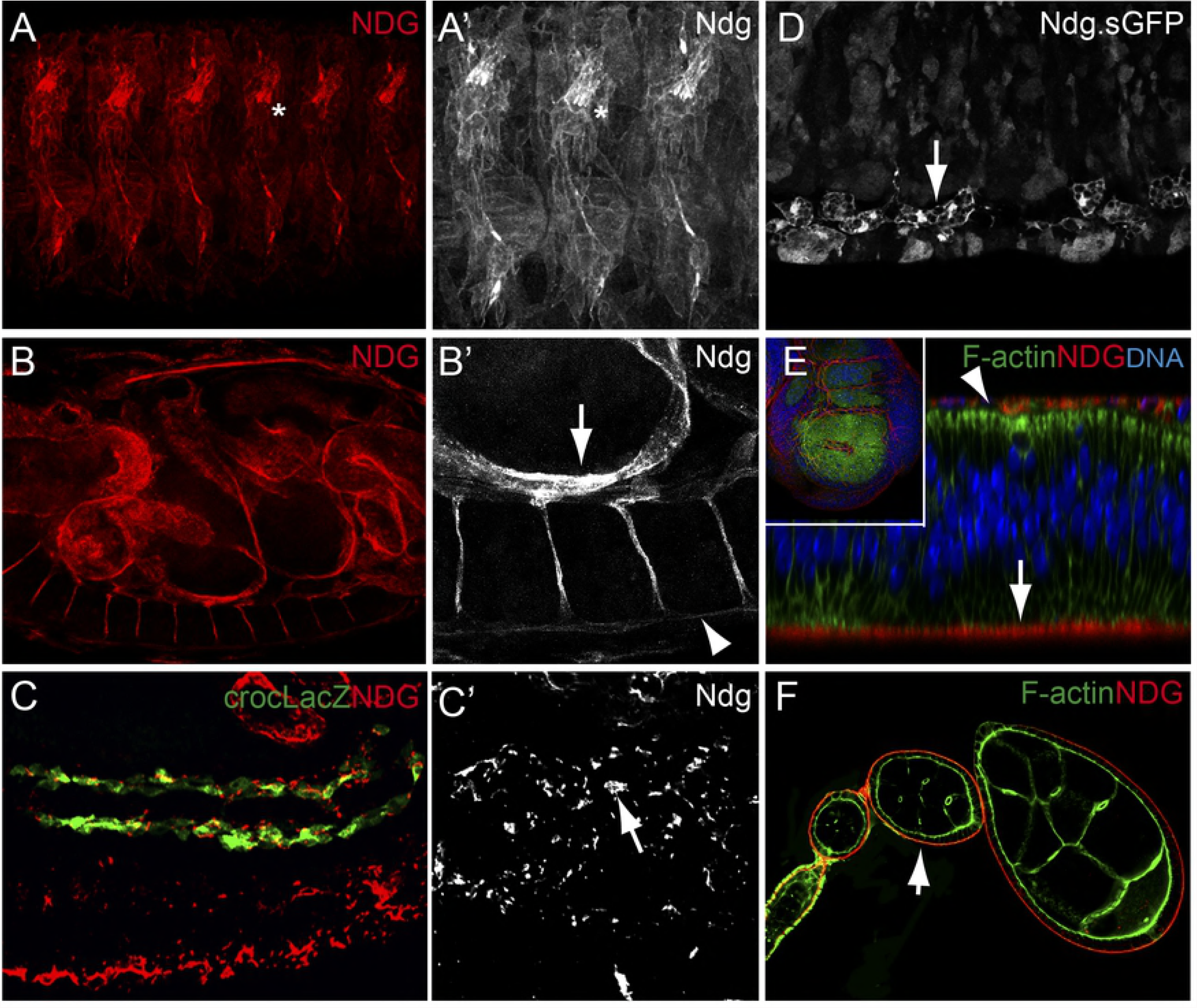
Distribution of the Ndg protein. (A-F) Confocal images showing embryonic (A-D), wing imaginal disc (E), and ovarian tissues (F) stained with anti-Ndg (A-C, and E-F) or anti-GFP (Ndg.sGFP, D). (A, B) In stage 16 wild-type embryos, Ndg (red) is found in the BMs surrounding most tissues, including muscles (A, A’), gut (B, B’ (arrow) and VNC (B, B’, arrowhead) and in chordotonal organs (asterisk). (C, C’, asterisk) Lateral view of a stage 13 embryo showing Ndg (red) accumulation around caudal visceral mesodermal cells visualized with the marker crocLacZ (green, arrow in C’). (D) Ndg is found in embryonic macrophages (arrow). (E) Ndg (red) is found at the basal surface of wing imaginal disc epithelial cells (arrow) and cells of the peripodial membrane (arrowhead). (F) Ndg (red) accumulates in the basal membrane around the follicular epithelium (arrow).

In addition, and similar to the other BM components, Ndg was found in the BMs that surround most larval tissues, including fat body, imaginal discs, tracheae, salivary glands, midgut, mature muscles and heart (Fig 1E and Fig 2), as well as in the follicular epithelium of the adult ovary (Fig 1F).

**Fig 2.**
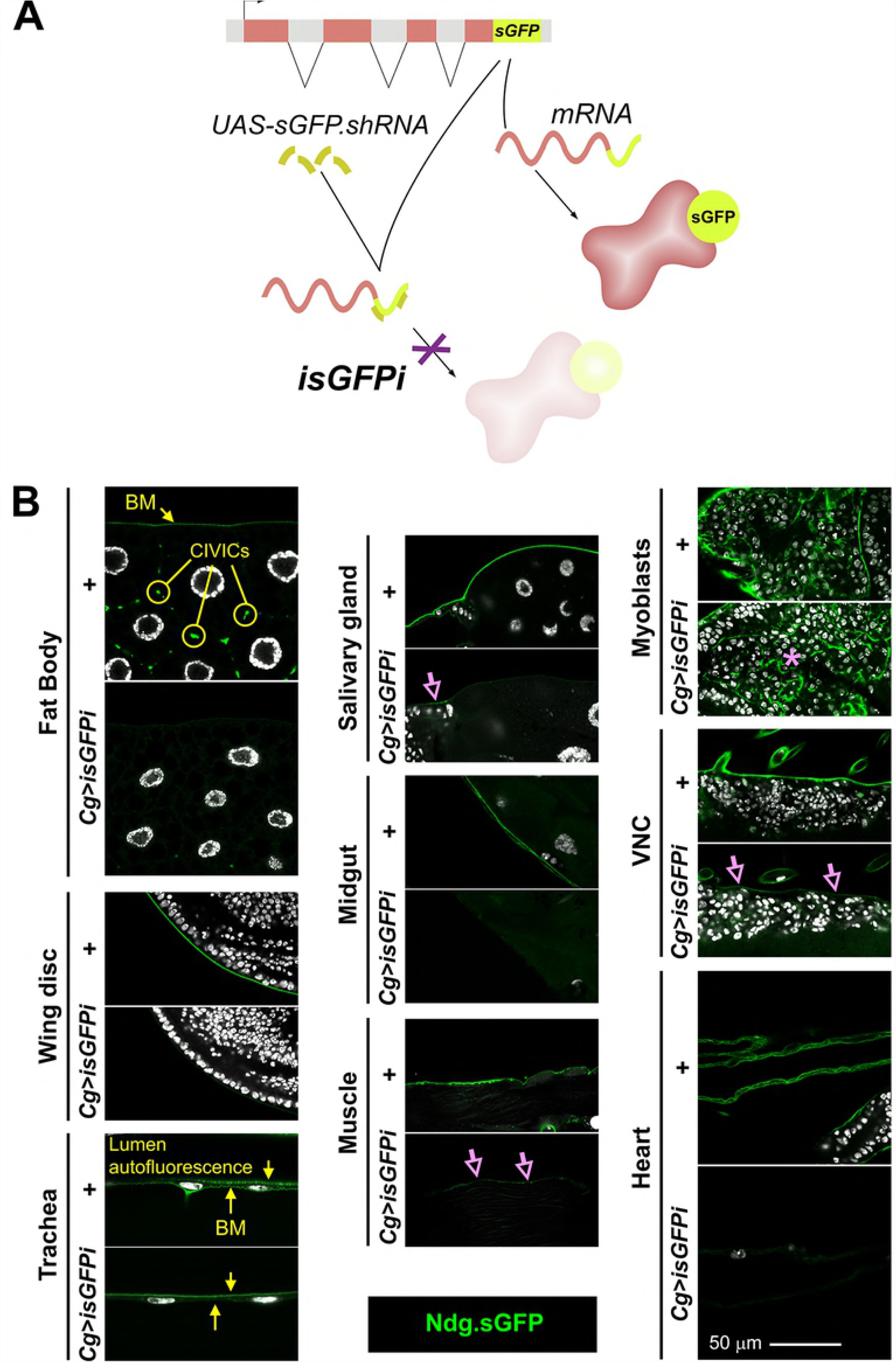
Fat body adipocytes and blood cells are the main source of Ndg in larval BMs. (A) Schematic representation of the in vivo sGFP interference approach (isGFPi). (B) Confocal images showing the localization of functional Ndg.sGFP fusion protein expressed under control of the endogenous promoter in different tissues of the third instar larva. Images compare tissues from control larvae (+) and larvae where Ndg.sGFP expression has been knocked down in fat body adipose tissue and blood cells (*Cg>isGFPi*). Ndg.sGFP signal is not observed in the BMs of *Cg> isGFPi* larvae, except in wing disc myoblasts (asterisk) and partially in the imaginal ring of the salivary gland, body wall muscles and VNC (purple arrows). Arrow and circles in the fat body panel indicate BM and CIVICs (Collagen IV Intercellular Concentrations), respectively. In tracheal images, apical cuticle autofluorescence is observed in the tracheal lumen (downward-pointing yellow arrow). Nuclei stained with DAPI (white).

### Fat body adipocytes and blood cells are the main source of Nidogen in larval BMs

A recent study has shown that macrophages are the major producers of BM components in the *Drosophila* embryo (32). To investigate the cellular origin of Nidogen in the developing fly, we designed a GAL4-driven UAS-controlled short hairpin against super-folder GFP (sGFP) to eliminate sGFP tagged Nidogen without disrupting normal function of endogenous untagged Nidogen (isGFPi, Fig 2A). This approach is similar to iGFPi (in vivo GFP interference, Pastor-Pareja and Xu, 2011) and iYFPi (Zang et al., 2015), which we previously used to show that fat body adipocytes are the major source of Collagen IV and Perlecan in the larva. We found here that isGFPi knock down of Ndg.sGFP driven by Cg-GAL4, which drives expression in fat body and blood cells (*Cg>isGFPi*), reduced the presence of Ndg.sGFP in the whole animal. Ndg.sGFP signal was largely reduced or undetectable in most tissues, including fat body itself, imaginal discs, tracheae, midgut and heart (Fig 2B). Deposition of Ndg.sGFP was only partially reduced in the ventral nerve cord, the imaginal ring of the salivary gland and body wall muscles, and was not affected in myoblasts, suggesting that these tissues could produce their own Ndg (Fig 2B). These results strongly suggest that, as it is the case for Collagen IV, fat body and blood cells are the main source of Ndg in the larvae.

We next decided to assess the origin of other components by performing the same essay for sGFP tagged Laminin, GFP tagged Col IV and YFP-tagged Perlecan (S1 Fig). We found that, similar to Nidogen, fat body and blood cells are the main source of the laminins, Col IV and Perlecan found in the BM of all tissues, except myoblasts and partially the VNC (S1 Fig). Tracheal cells seem also able to produce some Perlecan but not laminins or Col IV (S1 Fig). Finally, we found that wing imaginal disc cells and muscles can also produce their own laminins (S1 Fig). In all, our results show that although the four core BM components are largely produced by fat body adipocytes and blood cells, there are tissue-specific and component-specific exceptions to this rule.

### Generation of Nidogen mutant alleles

The *Drosophila* genome contains a single *Ndg* gene. To analyse *Ndg* requirements during development, we aimed to isolate a deficiency uncovering the gene *Ndg* (Fig 3A; see Materials and Methods). A series of deficiencies were generated by P-element imprecise excision of the transposon Mi{ET1}Ndg[MB12298] (S2 Fig, see Materials and Methods). All deficiencies eliminated the 5’ UTR and the first exon of the *Ndg* gene and at least two adjacent genes *Obp46a* and *CG12909* (S2 Fig). These deficiencies were all homozygous lethal.

**Fig 3.**
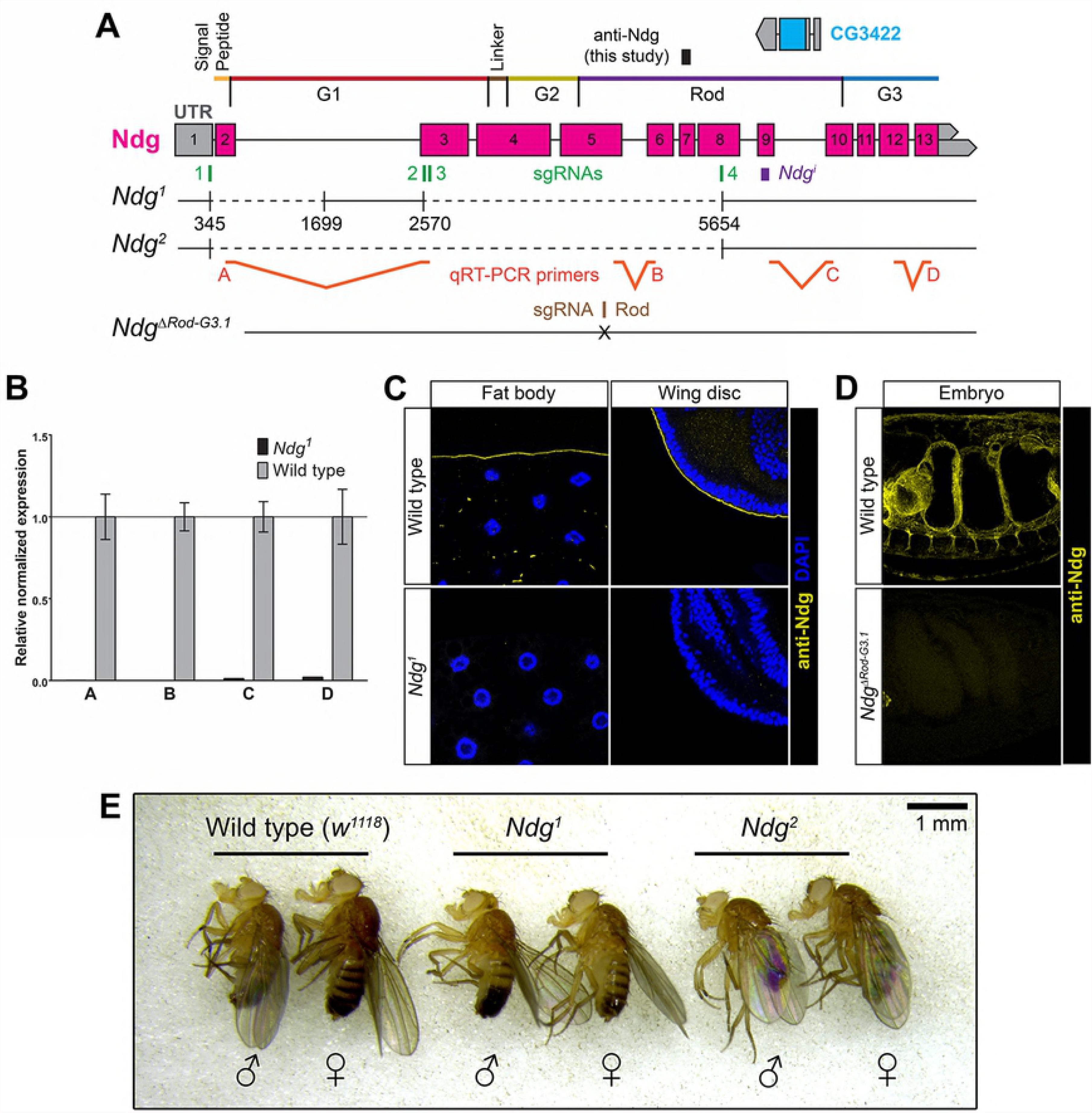
CRISPR/Cas9-generated Nidogen mutants are viable. (A) Schematic representation of the *Ndg* locus (2^nd^ chromosome), *Ndg* mutants generated, sgRNAs used for generation of mutants (1-4), sequence targeted by Ndg RNAi construct (*Ndg^i^*), epitope recognized by the rabbit anti-Ndg antibody generated in this study (black box) and qRTPCR primers used to molecularly characterized the *Ndg^1^* mutant (A-D). (B) Expression of *Ndg* mRNA in wild type (*w^1118^*) and *Ndg^1^* homozygous mutant larvae, assessed by qRT-PCR. Error bars represent 95% confidence intervals from three repeats. (C, D) Confocal images of larval fat body (C, left panels), wing imaginal discs (C, right panels) and embryos (D) from wild type (upper panels), *Ndg^1^* (C, lower panels) and *Ndg^ΔRodG3.1^* (D, lower panels) mutant animals stained with anti-Ndg antibody (yellow). (C) Nuclei stained with DAPI (blue). (E) Homozygous *Ndg^1^* and *Ndg^2^* mutants are viable and show no obvious morphological abnormalities.

As these deficiencies removed other genes, we took advantage of the CRISPR-Cas9 technology to isolate a series of specific *Ndg* alleles (Fig 3A; see Materials and methods). To generate *Ndg* null alleles, embryos were injected with Cas9 mRNA and a combination of four sgRNAs designed against the 5’UTR exon, exon 3 (two sg RNAs) and exon 8. Two mutant lines in which the intervening Ndg sequence had been deleted partially (*Ndg^1^*) or completely (*Ndg^2^*) (Fig 3A) were isolated. Gene CG3422, contained between exons 9 and 10 of the Ndg gene was not perturbed. Both mutations are predicted to be *Ndg* null alleles because of absence transcription start site. In fact, qRT-PCR, using different primers along the *Ndg* gene, showed no mRNA expression in *Ndg^1^* homozygous mutant larvae compared to wild type controls (Fig 3B). Furthermore, consistent with *Ndg^1^* and *Ndg^2^* being null alleles staining with our Ndg antibody could not detect presence of the protein in larval or embryonic tissues (Fig 3C and D).

As domain G3 has been postulated to be critical for binding of Ndg to laminins, we also isolated *Ndg* mutant alleles in which this domain and the adjacent rod domain were eliminated (ΔRod-G3 alleles) in order to analyze its function in the context of the whole organism. In this case, transgenic lines stably expressing an sgRNAs against exon 5 were generated and crossed to flies expressing Cas9 (see Materials and Methods). Three mutants alleles, *Ndg*^ΔRod-G3.1^, *Ndg*^ΔRod^G3.2 and *Ndg*^ΔRod-G3.3^, were selected (Fig 3A). Two of them, *Ndg*^ΔRod-G3.1^, *Ndg*^ΔRod-G3.2^, were deletions of five and eight base pairs that resulted in frame-shifts generating stop codons eight and seven amino acids after the shift, respectively. In the other one, *Ndg*^ΔRod-G3.3^, six base pairs were replaced by seven different ones, generating a frame-shift and a stop codon right after the shift. As expected, no staining using the antibody generated in this study was detected in *Ndg*^Δ*Rod-G3.1*^ homozygous embryos (Fig 3D and S3 Fig).

All CRISPR/Cas9 *Ndg* mutant alleles we generated were homozygous viable with no obvious morphological abnormalities (Fig 3E and data not shown). These data shows that, in contrast to double *Nid1 Nid2* knock out mice and similar to *C. elegans Nid-1* mutants, *Ndg* is dispensable for viability in *Drosophila*. In addition, alike in *C elegans*, elimination of *Ndg* results in reduced fertility in flies (S4A Fig). However, and in contrast to the phenotype observed when removing laminins, Col IV or perlecan (46) (24), no defects were observed in the shape of eggs laid by *Ndg* mutant females (S4B Fig).

### Nidogen is required for integrity of the BM of the larval fat body and adult flight muscles

Once shown that *Ndg* mutant flies are viable, we decided to analyze the effects of *Ndg* loss in the BMs of the fly. We could not detect defects in most of the BMs we analyzed, including those present in the embryo, larval epidermis, imaginal discs salivary glands, gut, muscles, ventral nerve cord and the follicular epithelium in the ovary. However, we did observe a clear defect in the BM surrounding the larval fat body (Fig 4A). The larval fat body is an organ formed by large polyploid cells (adipocytes) covered by a BM that separates it from the hemolymph (47). This BM contains, besides Ndg, the other three major components of BMs, Col IV, Laminins and Perlecan. Using tagged versions of these proteins, we found that the BM surrounding the fat body adipose tissue of *Ndg^1^* mutant larvae showed many holes, in contrast to the continuous appearance of the BM in wild type controls (Fig 4A). This phenotype was also observed when knocking down *Ndg* expression using the Cg-GAL4 (*Ndg^i^*, Fig 4A). BM rupture upon *Ndg* loss was also observed in transheterozygotes *Ndg^1^*/*Ndg^2^* and *Ndg^1^*/Df(2R)BSC281 (Fig 4B), confirming a requirement for *Ndg* to preserve fat body BM integrity. Furthermore, this phenotype was also observed in transheterozygotes *Ndg^1^*/ *Ndg*^ΔRod-G3.1^ larvae and in homozygous *Ndg*^ΔRod-G3.1^ (Fig 4B and data not shown), indicating a strong requirement of the Rod and G3 domains for this *Ndg* function. Confirming that these phenotypes reflected a loss of *Ndg* function, the Ndg.sGFP transgene rescued the rupture of fat body BMs in *Ndg^1^* mutants (Fig 4C).

**Fig 4.**
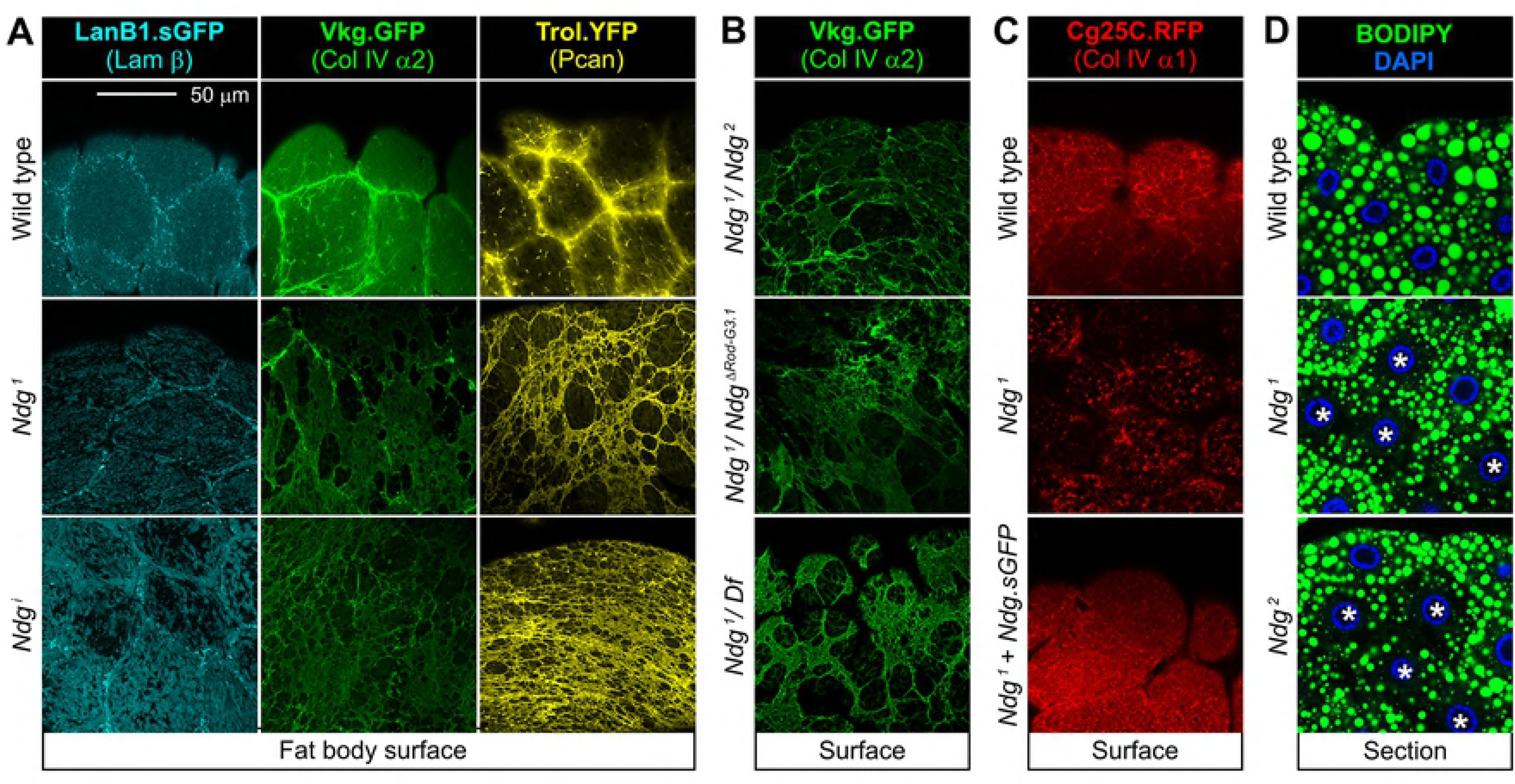
*Ndg* mutants show rupture of adipose tissue BMs. (A) Confocal images of the larvae fat body BM showing localization of Laminin (LanB1.sGFP, cyan), Collagen IV (Vkg.GFP, green) and Perlecan (Trol.YFP) from control (upper panels), *Ndg^1^* mutant (middle panels) and larvae where *Ndg* has been knocked down under control of CgGAL4 (*Ndg^i^*, lower panels). Loss of *Ndg* causes rupture of adipose tissue BMs. (B) Ruptured BMs (Vkg.GFP, green) in the larval fat body of transheterozygotes *Ndg^1^/Ndg^2^* (upper panel), *Ndg1*/ *Ndg^ΔRod-G3.1^* (middle panel) and *Ndg^1^/Ndg^Df(2R)BSC281^* (lower panel). (C) Fat body BM in wild type, *Ndg^1^* mutant and *Ndg^1^* mutant rescued with Ndg.sGFP. (D) Lipid droplets (neutral lipid dye BODIPY, green) in fat body of wild type (upper panel), *Ndg^1^* mutant (middle panel) and *Ndg^2^* mutant (lower panel) larvae. Asterisks point to cells with reduced content of lipid droplets. Nuclei stained with DAPI (blue).

We next investigated whether adipose tissue physiology was affected in *Ndg* mutants. To do this, we stained fat body adipocytes with neutral lipid dye BODIPY and found that the lipid content in *Ndg^1^* and *Ndg^2^* mutant adipocytes was reduced, with some cells presenting fewer and smaller lipid droplets than controls (Fig 4C).

In addition to fat body BM defects observed in *Ndg^1^* mutant larvae, we further discovered BM integrity defects in the flight muscles of the notum in *Ndg* mutant flies (S4C Fig). In addition, while flies appeared to fly normally and negative geotaxis climbing assays did not show differences with the wild type (not shown), Chill-Coma Recovery Time (CCRT) assays (48) showed increased recovery times after cold exposure in flies lacking *Ndg*, suggesting mild behavioral or motor defects (S4D Fig).

In summary, our results show that although Ndg is not critically essential for fly development and assembly of most BMs, it is necessary for the integrity of BMs around specific tissues, such as the larval fat body and the adult flight muscles, and for appropriate fertility and fitness of the fly.

### Functional analysis of the different Nidogen domains

In order to understand the function of Ndg and the basis for its specific requirement in the fat body, we performed a functional analysis of the different Ndg domains. To do that, we generated transgenic flies capable of expressing GFP tagged versions of the wild type Ndg protein as well as a series of mutant variants lacking one or several of these domains (Fig 5A).

**Fig 5.**
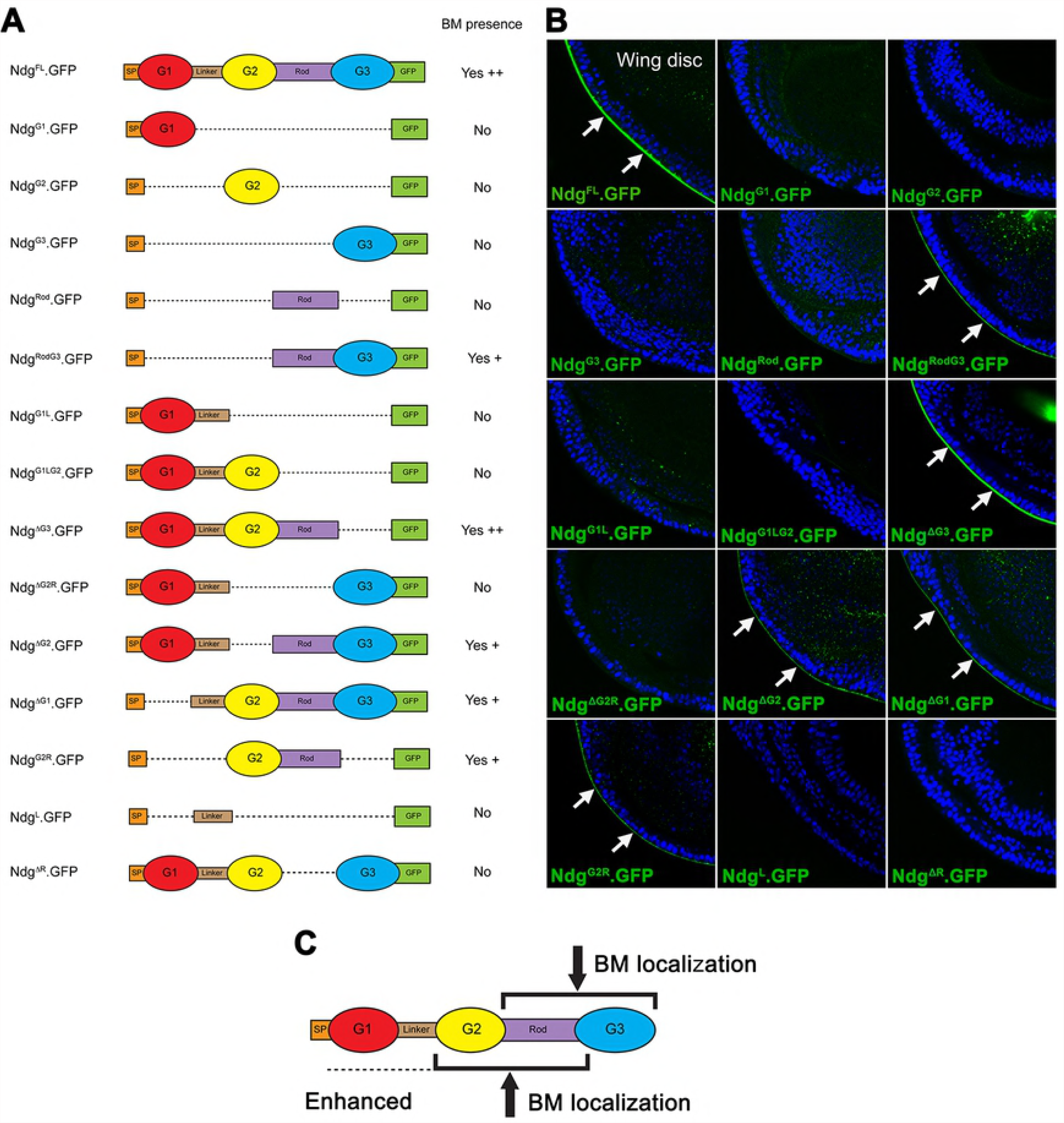
Rod domain is necessary but not sufficient for Nidogen localization to BMs. (A) Schematic depiction of Ndg full length (FL) and different Ndg deletions tested in this study. All Ndg versions are tagged with GFP in the C-terminal. SP=Signal Peptide. G1/2/3=Globular domains 1/2/3. (B) Confocal images of wing discs from larvae expressing the indicated versions of Ndg under control of Cg-GAL4. Correct BM localization, detected by GFP signal (green, arrows), is observed only for Ndg versions containing the Rod domain (Ndg^FL^, Ndg^RodG3,^ Ndg^ΔG3^, Ndg^ΔG2^, Ndg^ΔG1^ and Ndg^G2Rod^) but neither Ndg^Rod^ nor any other single domain can localize to the BM by itself. Nuclei stained with DAPI (blue). (C) Model of domain requirements for proper Ndg localization to BMs.

First, we addressed whether these proteins localized normally to BMs. When expressed in the fat body and blood cells under control of the Cg-GAL4 driver, full length Ndg (Ndg^FL^.GFP) was able to localize to the BMs of imaginal discs, as expected (Fig 5B). A similar analysis of the localization properties of the deletion constructs showed that no single domain of the protein was capable by itself to confer localization to BMs, suggesting cooperative interactions among domains are required for BM localization (Fig 5B). In addition, analysis of the localization of proteins in which a single domain was deleted (Ndg^ΔG1^, Ndg^ΔG2^, Ndg^ΔG3^ and Ndg^ΔRod^) showed that the only domain absolutely required for BM localization was the Rod domain (Fig 5B). However, the Rod domain was insufficient to drive protein localization on its own (Ndg^Rod^), but required the presence of the G2 or the G3 domains (Ndg^G2Rod^ and Ndg^RodG3^, respectively, Fig 5B). This result is supported by our analysis of the localization of Ndg in our *Ndg*^ΔRod-G3^ mutant embryos using an antibody raised against the G2 domain of Ndg (49) (S3 Fig). Staining with this antibody showed that the mutant protein Ndg^ΔRod-G3^ did not localize to embryonic BMs, but it was still present in the chordotonal organs (S3 Fig). This result suggests that the truncated protein lacking the Rod and G3 domains is expressed by chordotonal organs and other tissues, but is not deposited into BMs. Altogether, these results show that the Rod domain is required but not sufficient for Ndg BM localization. They also show that G2+Rod and Rod+G3 are minimal alternative units capable of conferring BM localization to the Ndg protein, with G1 enhancing G2+Rod dependent-localization.

As mentioned in the introduction, analysis of the binding properties of Ndg domains and crystal structure have suggested that the G3 domain binds to Laminin, whereas the G2 domain binds to Col IV, with no clear function ascribed so far to the G1 domain. To investigate this in vivo, we assayed the localization abilities of the Ndg^ΔG1^, Ndg^ΔG2^ and Ndg^ΔG3^ mutant proteins in the BMs of fat bodies where the expression of either laminins or Col IV had been knocked down. We found that knock-down of *LanA* or *Cg25C*, encoding a Laminin a chain and Col IV α1, caused a marked reduction in the incorporation of full length Ndg^FL^.GFP (Fig 6A-C). In addition, while knocking down *LanA* had not effect on Ndg^ΔG3^ localization, *LanA* loss caused a marked reduction in the localization of both Ndg^ΔG1^ and Ndg^ΔG2^ (Fig 6A and B). Conversely, knocking down *Cg25C* resulted in a strong reduction in Ndg^ΔG3^ localization and significant but not as drastic effects on the localization of Ndg^ΔG1^and Ndg^ΔG2^ (Fig 6A and C). Altogether, these results show that localization directed by the G3 domain depends on laminins, whereas localization by the G1 and G2 domains depends on Col IV.

**Fig 6.**
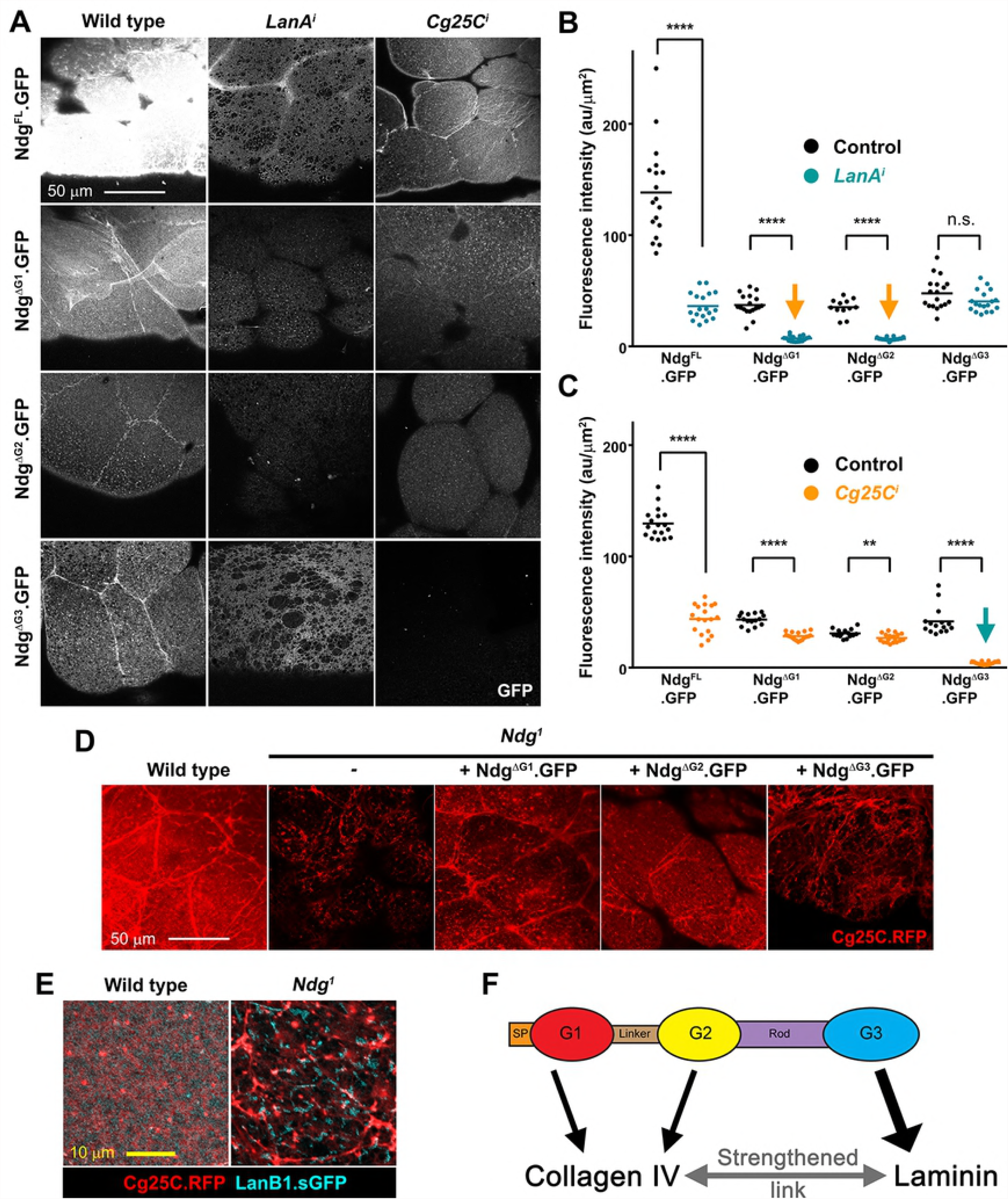
The G3 domain is essential for Ndg function. (A) Confocal images showing localization in fat body BM of the indicated GFP-tagged versions of Ndg, detected by GFP signal (white), in wild type (right column) and upon knock down of Laminin (*LanA^i^*, middle column) and Collagen IV (*Cg25C^i^*, left column). (B, C) Quantification of GFP signal intensity of indicated GFP-tagged Ndg versions in the BM of wild type control, *LanA^i^* (B) and *Cg25C^i^* (C) fat body. Each dot represents a single measurement of intensity inside a 500μm^2^ square. Horizontal lines indicate the mean value (****: p<0.0001). (D) Confocal images of the larvae fat body BM (Cg25C.RFP in red) in control, Ndg1 mutant, and *Ndg^1^* mutant expressing Ndg^ΔG1^, Ndg^ΔG2^ or Ndg^ΔG3^. Integrity of the BM is restored by Ndg^ΔG1^ and Ndg^ΔG2^ but not Ndg^ΔG3^. (E) Confocal images showing uncoupling of Collagen IV (Cg25C.RFP, red) and Laminin (LanB1.sGFP, cyan) in *Ndg^1^* mutant fat body (right panel) compared to control (left panel). (F) Model for the role of the G1, G2 and G3 domains on Ndg binding to Laminin and Collagen IV.

Next we tested the ability of the Ndg mutant proteins lacking the G1, G2 or G3 domains, all three capable of localizing to BMs, to rescue the fat body BM defects observed in *Ndg^1^* mutant larvae (Fig 6D). Overexpression of the mutant variants Ndg^ΔG1^ or Ndg^ΔG2^ was able to rescue integrity of the fat body BM (Fig 6D), as imaged with Cg25C.RFP (50) (50). In contrast, expression of the mutant form Ndg^ΔG3^ failed to rescue BM rupture, indicating that G3 is a key domain for Ndg function, while the G1 and G2 domains may function in a partially redundant way. This is supported by our results showing that *Ndg^1^*/*Ndg*^ΔRod-G3.1^ transheterozygous mutant larvae show fat body BM defects indistinguishable from those found in *Ndg^1^* homozygotes (Fig 4A and B).

The localization and rescue properties of the different domains of Ndg suggest that Ndg may indeed act as a linker between Laminin and Collagen IV, as originally proposed. Confirming this, simultaneous imaging of Collagen IV and Laminin in fat body BMs shows that in the *Ndg^1^* mutant Laminin and Collagen IV appear separate from each other when the damaged BM is observed at high magnification (Fig 6E). In all, these results are consistent with a function of Ndg as a linker of the Col IV and Laminin networks (Fig 6F). This linker function would depend on binding to Laminin through G3 and to Col IV through either G1 or G2.

### Role of Laminins, Collagen IV and Perlecan in Nidogen incorporation into BMs

We have previously shown that *Drosophila* laminins are critical for proper assembly of other ECM components in the BM of embryonic tissues (35). Furthermore, recent studies have shown that there is a temporal hierarchy of BM components expression in the *Drosophila* embryo, with laminins being expressed first, followed by Col IV and then Perlecan (32). This seems to be critical for proper formation of the BM around the embryonic VNC. Thus, while elimination of laminins affects both Col IV and Perlecan deposition, laminin incorporates in the absence of any of these two components and Perlecan requires Col IV (32). The requirements of these BM proteins for Ndg incorporation into embryonic BMs are still unknown. Here, we decided to investigate this by analysing Ndg expression in embryos devoid of the other BM components. We found that depletion of LanB1 results in a strong reduction of Ndg accumulation in the gut, muscles and VNC (Fig 7A and B). However, elimination of SPARC, a carrier for Col IV (31) (51), or Perlecan did not prevent Ndg deposition into embryonic BMs (Fig 7C and D). This is in agreement to what we have previously found in wing imaginal discs where Col IV is not required for Ndg localization (31).

**Fig 7.**
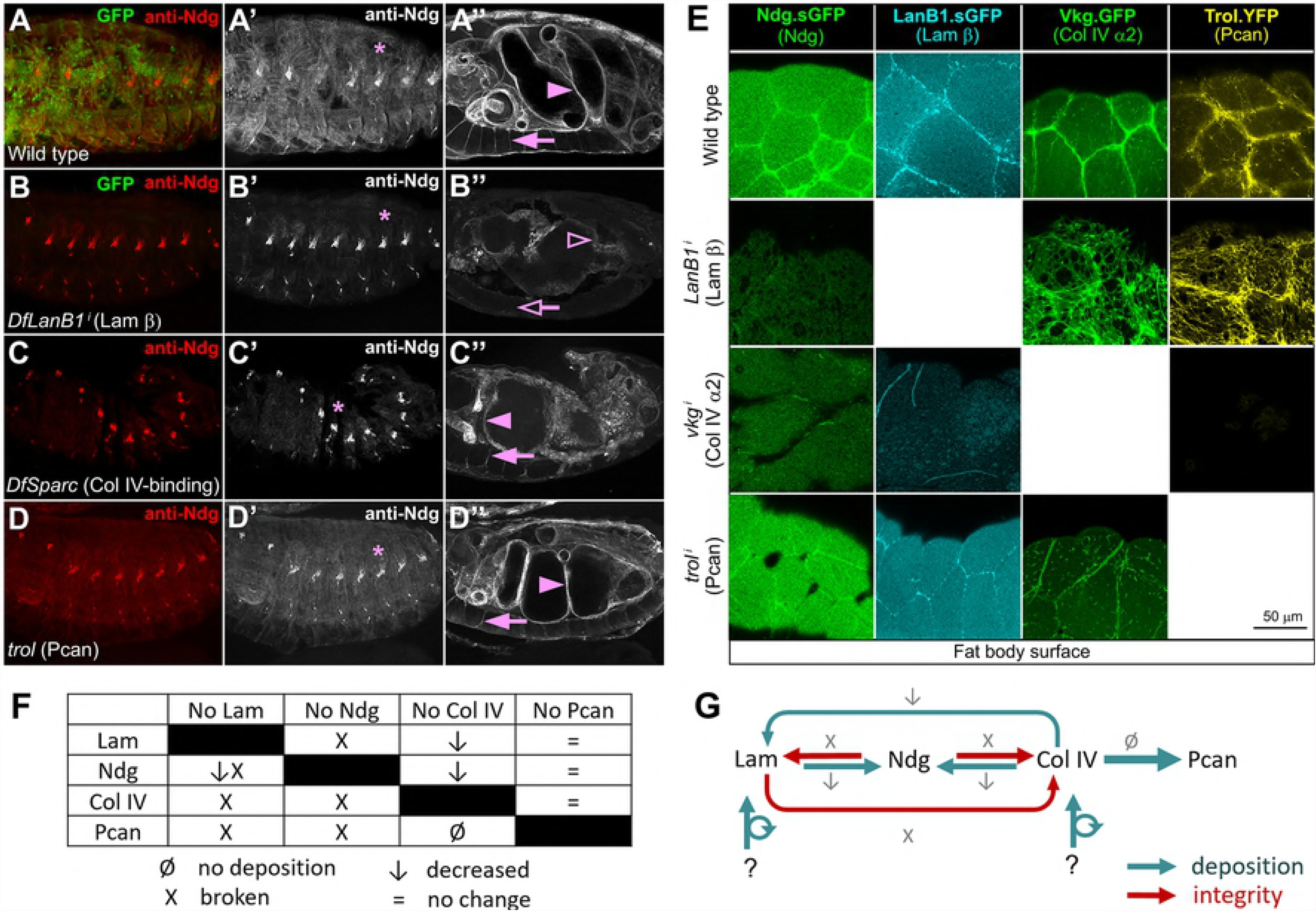
Laminins and Col IV are required for proper Ndg incorporation into adipose tissue BMs. (A-D”) Confocal images showing stage 16 embryos stained with anti-Ndg (red) and anti-GFP (green). Images compare control embryos (A) with embryos depleted of the different BM components (B-D). (A-A”) In control embryos, Ndg localizes to the BM of muscles (A’), gut (A”, arrowhead) and VNC (A”, arrow) and in chordotonal organs (A’, asterisk). (B) In Laminin depleted embryos Ndg is strongly reduced in the BM of muscles (B’), gut (B”, hollow arrowhead) and VNC (B”, hollow arrow) and not affected in chordotonal organs (B’, asterisk). (C-D) In contrast, Ndg deposition in these embryonic BMs is not affected in either Sparc (C-C”) or Perlecan (D-D”) mutant embryos. (E) Confocal images of the fat body BM showing localization of Ndg (Ndg.sGFP, green), Laminin (LanB1.sGFP, cyan), Collagen IV (Vkg.GFP, green) and Perlecan (Trol.YFP). Images show larvae fat body from wild type (upper panels) and larvae where *LanB1, vkg or trol* have been knocked down using the Cg-GAL4 (lower panels). (F) Table summarizing effects of the absence of each of the four major BM components on the other components. (G) Model for the mutual relations of Laminin, Nidogen, Collagen IV and Perlecan in the BM of the fat body.

Next, we tested the requirements of laminins, Col IV and Perlecan for Ndg incorporation into the BM of the larval fat body. To this end, we analysed the expression of the transgene Ndg.sGFP in the fat body of larvae where we had knocked down expression of BM components under the control of the Cg-Gal4 driver. We found that the knock down of laminins or Col IV, but not of Perlecan, caused a reduction in the amount of *Ndg* in fat body BMs (Fig 7 and S5 Fig), consistent with our functional analysis of the different Ndg domains (Fig. 6).

We aditionally decided to analyze the mutual requirements of the remaining components of the adipose tissue BM. We found that loss of Col IV result in a strong reduction in Laminins levels and in a depletion of Perlecan (Fig 7E and S5 Fig). This is in agreement with previous results showing that knocking down *vkg* with *hsp70-gal4*, which is a heat shock inducible promoter, reduced the presence of Nidogen and Laminin in fat body BM (52). In contrast, similar to the loss of Ndg (Fig 4), absence of laminins led to rupture of the BM without apparent reduction in collagen or Perlecan levels (Fig 7B and S5 Fig). Finally, knock down of Perlecan did not affect the presence of any of the other components, consistent with the notion that it is a terminal BM component (Fig 7E) (31).

In summary, these results show that Ndg incorporation into embryonic and fat body BMs depends on both Laminins and Collagen IV. They also suggest a model for the assembly and maintenance of the adipose tissue BM in which Ndg is not not essential for the incorporation of other components, but reinforces the connection between laminins and collagens, thus preventing the rupture of the BM (Fig 7G).

### Genetic interactions unmask a wider involvement of Nidogen in BM stability

To finally ascertain whether Nidogen incorporation had a wider stabilizing role on BMs despite limited phenotypic defects in the mutants, we tested genetic interactions with other genetic conditions compromising BM functionality. *LanA^216^* and *LanA^160^* are two embryonic lethal mutant alleles of *LanA* (53). While *LanA^216^/LanA^160^* 3^rd^ instar larvae showed an elongated ventral nerve cord (VNC), no defects in VNC condensation were observed in *LanA^216^/+*, *LanA^160^/+* or *Ndg^1^*. In contrast, we found that *Ndg^1^* mutants heterozygous for *LanA^216^/+* or *LanA^160^/+* showed VNCs that are significantly more elongated than those found in *LanA^216^/+* or *LanA^160^/+*, respectively (Fig 8A and B). In addition, we found that Ndg interacted genetically with Perlecan. Thus, while single knock down of either Ndg or Perlecan in the whole fly, using actin-GAL4, produced normal-looking pupae and viable adults, the double knock out of these genes caused a decrease in the size of pupae, which were unable to develop to adulthood (Fig 8). The combined interaction was exacerbated when knock down was driven at 30°C, a temperature at which GAL4-driven transgene expression is higher (54) (55). In summary, these results prove that Nidogen interacts genetically with Laminins and Perlecan, suggesting a more general role of Nidogen in maintaining BM stability and consistent with its remarkable evolutionary conservation.

**Fig 8.**
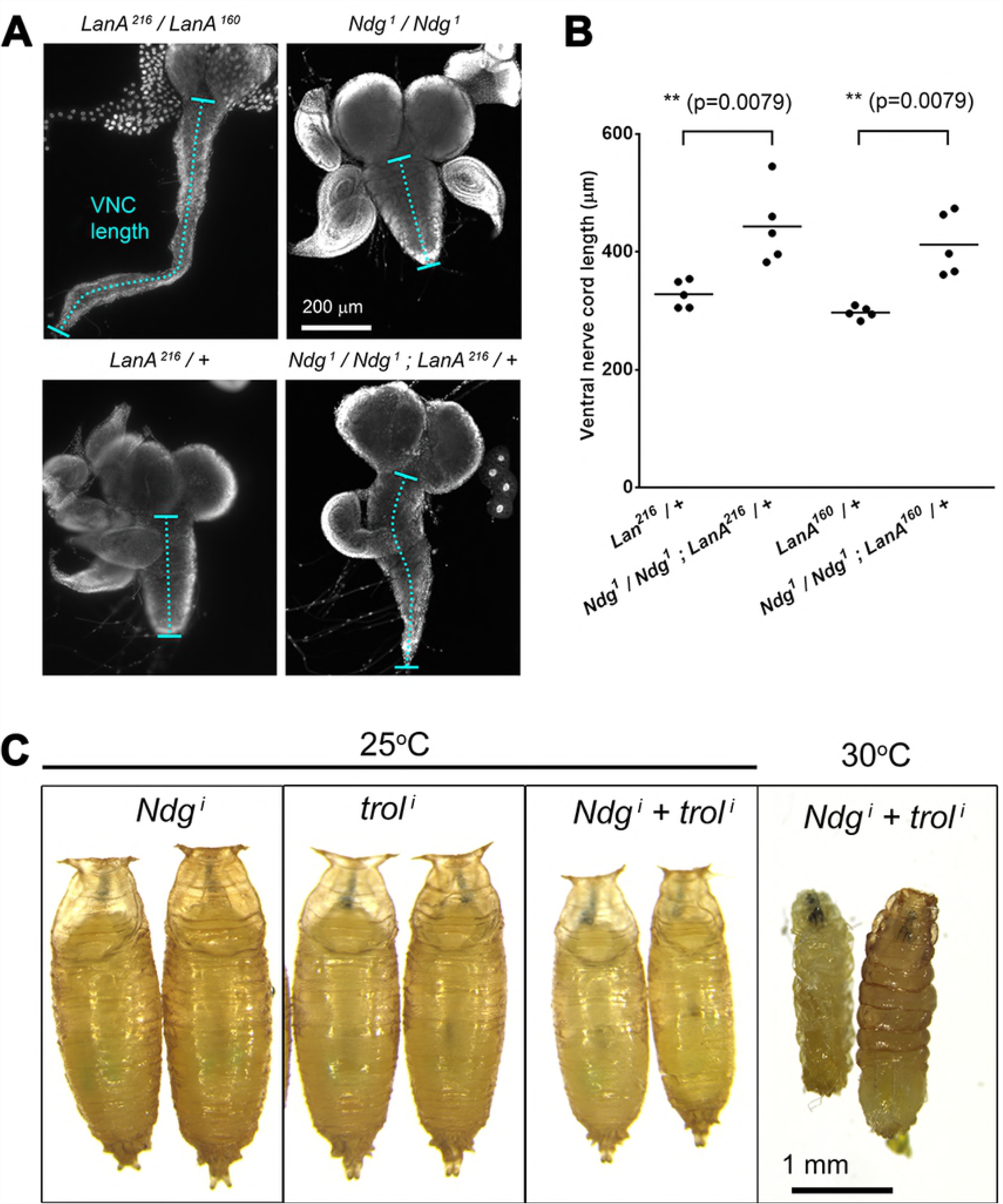
Ndg loss enhances Laminin and Perlecan loss phenotypes. (A) Larval ventral nerve cord (VNC) in *LanA^216^/LanA^160^* (upper left panel), *Ndg^1^/Ndg^1^* (upper right panel), *LanA^216^/+* (lower left panel) and *Ndg^1^/Ndg^1^; LanA^216^/+* (lower right panel) larvae.(B) Quantification of VNC length in *LanA^216^/+*, *Ndg^1^/Ndg; LanA^216^/+*, *LanA^160^/+* and *Ndg^1^/Ndg; LanA^160^/+*. Each dot represents an individual measurement. Horizontal lines indicate the mean value. Differences with the control were significant in non-parametric Mann-Whitney tests (**; p<0.01). (C) Images of pupae where *Ndg*, *trol* or both *Ndg* and *trol* have been knocked down under using act-GAL4 at either 25°C or 30°C.

## Discussion

BMs are thin extracellular matrices that play crucial roles in the development, function and maintenance of many organs and tissues (56). Critical for the assembly and function of BMs is the interaction between their major components, Col IV, laminins, proteoglycans and Ndg (57). Both the ability of Ndg to bind laminin and Col IV IV networks and the crucial requirements for Laminins and Col IV in embryonic development (58, 59) anticipated a key role for Ndg during morphogenesis. However, experiments showing that elimination of Ndg in mice and *C. elegans* is compatible with survival cast doubt upon the crucial role for Ndg in organogenesis as a linker of the crucial Laminin and Col IV networks within the BM. Here, we have isolated mutations in the single *Drosophila Ndg* gene and found that, as it is the case in mammals and *C. elegans*, *Ndg* is not generally required for BM assembly and viability. However, *Ndg* mutant flies display mild motor or behavioral defects. In addition, similar to mammals, we show that the nidogen-deficient flies show BM defects only in certain organs, suggesting tissue-specific roles for *Ndg* in BM assembly and maintenance. Finally, our functional study of the different Ndg domains challenges the significance of some interactions derived from in vitro experiments while confirming others and additionally revealing a new key requirements for the Rod domain in Ndg function and incorporation into BMs.

Results from cell culture and in vitro experiments have led to propose a crucial role for Ndg in BM assembly and stabilization. Recombinant Ndg promotes the formation of ternary complexes among BM components (60). In addition, incubation with recombinant Ndg or antibodies interfering with the ability of Ndg to bind laminins results in defects in BM formation and epithelial morphogenesis in cultured embryonic lung, submandibular glands and kidney (22, 23). However, elimination of Ndg in model organisms has shown that Ndg is not essential for BM formation per se but required for its maintenance in some tissues. Thus, while the early development of heart, lung and kidney, prior to E14, is not affected in nidogen-deficient mice, defects in deposition of ECM components and BM morphology were observed at E18.5 (11). In addition, whereas BM components localized normally underneath the apical ectodermal ridge (AER) in Nidogen-deficient mice at birth, this BM breaks down at later stages (12). In contrast, removal of *Ndg* does not impair assembly or maintenance of any BM in *C.elegans* (13). Here, we show that in *Drosophila*, as it is the case in mammals (11, 12), different BMs have different requirements for *Ndg*. Thus, while elimination of *Ndg* in *Drosophila* does not impair embryonic BM assembly or maintenance, it results in rupture of the BM in fat body and flight muscles. The basis for this tissue-specificity of Ndg requirements is currently unknown. Recent experiments have shown that there is a tissue-specific hierarchy of expression and incorporation of BM proteins in the *Drosophila* embryo, with Laminins being expressed first followed by Col IV and finally Perlecan (32). Laminins and Col IV can reconstitute polymers in vitro that resemble the networks seen in vivo (34) (61). In this context, Laminins and Collagens could self-assemble into networks in the embryo as they are produced, being this sufficient to assemble a BM capable to sustain embryonic development in the absence of the two subsequent components, Ndg and Perlecan. We also show here that, while fat body and blood cells are the source of the majority of the proteins in larval BMs, there are notable exceptions, a fact that highlights a diversity in the origins of BM components in different tissues. Thus, fat body produces entirely all its BM, the larval heart receives it all from the hemolymph, imaginal discs produce a portion of their laminins and similarly for tracheae with respect to Perlecan. These differences in the source of the components for the different tissues (incorporated vs. self-produced) may impose different assembly mechanisms, a possibility to study in more detail in the near future. In addition, although BM components are universally present in numerous tissues and organs, they are diverse depending on tissue and developmental stage (reviewed in (62). This heterogeneity arises from variations in protein subtypes, such as the two alternative Laminin α chains or the numerous Perlecan isoforms. Heterogeneity may also stem from differences in relative amounts of each component and posttranslational modifications thereof. In this respect, it is possible that BM assembly of the *Drosophila* fat body and adult flight muscles of the notum is such that is more dependent on *Ndg* function for its formation and stability than BMs found in other tissues. Finally, dynamics of BMs can orchestrate organ shape changes. Reciprocally, the associated tissues can control properties of BMs by, for instance, expressing a specific repertoire of ECM receptors or remodeling factors. In this context, it is also possible that fat body or adult flight muscles sculpt BMs with properties demanding a high requirement of *Ndg* function.

We find here that *Ndg* mutant flies are less fertile and behave differently with respect to wild type in Chill-Coma Recovery Time assays. The physiological mechanisms underlying the response in insects to critical thermal limits remain largely unresolved. The onset and recovery of chill coma have been attributed to defects in neuromuscular function due to depolarization of muscles fibre membrane potential (63). Interestingly, flight muscle fibre membrane is particularly strongly depolarized upon exposure to low temperatures in *Drosophila* (63). In this context, the defects we observed in the BM of adult flight muscles in the absence of *Ndg* could be behind the defective response of *Ndg* mutant flies to chill coma recovery assays. Finally, we have found in a preliminary analysis that *Ndg* mutant larva show altered immune response to microbial infection (data not shown). All together, these results show that although not critical for survival, Ndg is required for overall fitness of the fly.

All Nidogen proteins consist of three globular domains, G1 to G3, and two connecting segments, one Rod domain separating G2 and G3 and a flexible link between G1 and G2. Crystallographic and binding epitope analyses using recombinant proteins of the domains of the mouse nidogen-1 have demonstrated high affinity binding of domain G2 for Col IV and perlecan and domain G3 for the laminin γ1 chain and col IV, and no activity for the Rod domain (4-7, 64). In addition, recent physicochemical studies analyzing the solution behavior of full length purified nidogen-1 confirmed the formation of a high affinity complex between the G3 domain of nidogen-1 and the laminin γ1 chain, and excluded cooperativity effects engaging neighboring domains of both proteins (65). However, little is known about the functional meaning of the binding abilities of Ndg on its localization and function in BM assembly in vivo. In fact, mutant *C. elegans* animals carrying a deletion removing the entire G2 domain of NID-1 are viable and show no defects on Ndg or Col IV localization in BMs (13). These results demonstrate that besides the strong sequence conservation between *C. elegans* and mammalian G2 domains, NID-1 localization appears to occur independently of this domain. Here, we show that, as it is the case in *C. elegans*, the *Drosophila* G2 domain is not essential for either Ndg localization or function. A possible explanation for this result is that although some of the modules present in BM components are conserved, there are variations in sequence and structure that might be enough to confer binding specificity to the different proteins. For instance, the IG3 domain of the mouse Perlecan that binds to G2 β-barrel domain in Nidogen is strikingly conserved in all metazoans. However, this domain has not been identified in either *Drosophila* or *C. elegans* Perlecan (66) (67). This result suggests that either the Perlecans present in these organisms are too distant in evolution from the mouse proteins for these domains to be conserved or that Perlecans may only bind Nidogen in mammals. Previous studies aimed to characterize the biological significance of the nidogen-laminin interactions have targeted the nidogen-binding module of the laminin γ1 chain, showing that this domain is required for kidney and lung organogenesis (22) (68). However, the role of the Nidogen G3 domain has not yet been addressed directly. Here, we show that the G3 domain is essential for Ndg localization, supporting a role for nidogen-laminin interactions on Ndg function. In addition, in contrast to what has been shown in mammals (see above), our results unravel a key role for the Rod domain in Nidogen localization. Again, an explanation for this result could hinge on variations in the Nidogen between species. In fact, one of the major differences between *Drosophila* and mammalian Nidogens lies on the Rod domain. Thus, while vertebrates have four EGF repeats and one or two thyroglobulin repeats *Drosophila* and *C. elegans* have 12 and 11 EGF repeats, respectively. Alternatively, conclusions derived from in vitro studies may not always be applicable to the circumstances occurring in the living organism. Furthermore, the appearance of new in vitro studies combining different techniques has revealed the existence of multiple nidogen-1/laminin γ1 interfaces, which include, besides the known interaction sites, the Rod domain (64).

Different BM assembly models have been proposed over the last thirty years. Based upon biochemical studies and rotary shadow electronic microscopic visualization, the BM assembly model firstly proposed that Collagen IV self-assembles into an initial scaffold, followed by Laminin polymerization structure attachment mediated by Perlecan (69, 70). However, more recent studies have postulated a contradicting model for in vivo systems. The most widely endorsed model was that the polymer structure is initiated by Laminin scaffold built through self-interaction, bridged linked by Nidogen and Perlecan and finally completed by another independent network formed by Col IV self-interaction (4). Here, we studied in some detail the hierarchy of BM assembly in the *Drosophila* larval fat body. Thus, while the requirements for *Drosophila* laminins in the incorporation of other ECM components into BMs are preserved between tissues, this is not the case for Collagen IV. For instance, absence of Col IV does not completely prevent deposition of Laminin in the fat body, but remarkably reduces it; in contrast, no such drastic effect has been observed in wing discs or embryonic BMs (this study, Pastor-Pareja an Xu, 2011 and (32), suggesting that Collagen IV is not affecting Laminin incorporation in these other tissues to the same degree or that it does not affect it at all. In addition, we found that BM assembly in *Drosophila* also differs from that in mammals and *C. elegans*. In this case, the divergences may arise during evolution, when different organisms might have incorporated novel ways to assemble ECM proteins to serve new specialized functions.

Nidogen has been proposed to play a key role on BM assembly based on results from in vitro experiments and on its ability to serve as a bridge between the two most abundant molecules in BMs, laminins and type IV collagens. However, phenotypic analysis of its knock out in mice and *C. elegans* have called into question a general role for nidogens in BM formation and maintenance. Here, we show that although Ndg is dispensable for BM assembly and preservation in many tissues, it is absolutely required in others. These differences on Ndg requirements stress the need to analyze its function in vivo and in a tissue-specific context. In fact, we believe this should also be the case when analyzing the requirements of the other ECM components for proper BM assembly, as we show here they also differ between species and tissues. One has to be cautious when simplifying functions to the different BM proteins or their domains based on experiments performed in vitro or in a tissue-specific setting. This might be especially relevant when trying to apply the conclusions derived from these studies to our understanding of the pathogenic mechanisms of BM-associated diseases or to the development of innovative therapeutic approaches.

## Materials and Methods

### Fly strains

Standard husbandry methods and genetic methodologies were used to evaluate segregation of mutations and transgenes in the progeny of crosses (71). The following stocks were used:

The FTG, CTG and TTG balancer chromosomes, carrying twist-Gal4 UAS-2EGFP, were used to identify homozygous *Ndg^ΔRod-G3^* mutants (Halfon et al., 2002). For the generation of Ndg deficiencies the following stocks were used (all from Bloomington *Drosophila* Stock Center): *Mi{ET1}Ndg^MB12298^, w; BlmN1/TM3, Sb1* (Witsell et al., 2009), *w; Sco / Sm6aP(hsILMiT)2,4*, *w; Gla/CyO*, and *Df(2R)BSC281*.

*w; tub-GAL80^ts^*, *y v; UAS-trol.RNA ^TRiP.JF03376^, y v; Ndg.RNAi^TRiP.HMJ24142^*, *y v sc; UAS-LanB2.RNAi^TRiP.HMC04076^, y v; UAS-LanA.RNAi^TRiP.JF02908^* and *y v sc; UAS-EGFP.shRNA* (BDSC 41560) are from Bloomington *Drosophila* Stock Center. *w; Ndg.sGFP^fTRG.638^, w; LanB1.sGFP^fTRG.638^, w; UAS-LanB1.RNAi^VDRC.v2312^, w; UAS-trol.RNAi^VDRC.v24549^* and *w; UAS-Cg25C.RNAi^VDRC.v28369^* were obtained from Vienna Drosophila Resource Center. *y w; vkg^G454^. GFP* and *w trol^CPTI-002049^.YFP* were from Drosophila Genomics Resource Center. *w; UAS-vkg.RNAi^NIG.16858R-3^* was from National Institute of Genetics.

Other strains used were: *DfLanB1/CTG* (35), *Df (3R)BSC524/CTG* (72), *Trol^null^/FMZ* ((73)), *w; UAS-Cg25C.RFP3.1* (Zang et. al, 2015). *w; LanA^160^/TM6B*, *croc-lacZ* (44) and *w; LanA^216^/TM6B* are gifts from Luis Garcia-Alonso.

Lines generated in this study are: *w; sGFP^RNAi^.attP40*, *w; Ndg^1^, w; Ndg^2^, Ndg^ΔRod-G.3.1^*, *Ndg^ΔRodG.3.2^, Ndg^ΔRod-G.3.3^. w; UAS-Ndg^FL^. GFP*, *w; UAS-Ndg^G1^. GFP*, *w; UAS-Ndg^G2^. GFP*, *w; UAS-Ndg^G3^. GFP*, *w; UAS-Ndg^Rod^.GFP*, *w; UAS-Ndg^RodG3^. GFP*, *w; UAS-Ndg^G1L^. GFP*, *w; UAS-Ndg^G1LG2^. GFP*, *w; UAS-Ndg^ΔG3^. GFP*, *w; UAS-Ndg^ΔG2R^. GFP*, *w; UAS-Ndg^ΔG2^. GFP*, *w; UASNdg^ΔG1^. GFP*, *w; UAS-Ndg^G2R^.GFP*, *w; UAS-Ndg^L^. GFP* and *w; UAS-Ndg^ΔR^. GFP*.

The GAL4-UAS system was used to drive expression of transgenes and RNAi constructs in larval fat body and hemocytes (blood cells) under control of Cg-GAL4 (BDSC 7011) or BM-40-SPARC-GAL4 (gift from Hugo Bellen), and ubiquitously with act-GAL4.

For Collagen IV knock down experiments (*vkg^i^* and *Cg25C^i^*), thermosensitive GAL4 repressor GAL80^ts^ was used to prevent embryonic lethality. Cultures were grown at 18°C for 6 days, followed by transfer of cultures to 30°C (L2 stage) and dissection two days later (L3 stage).

### Transgenic flies

### sGFP RNAi

Short hairpin oligoes to knock down sGFP were designed following instructions in DSIR website (http://biodev.extra.cea.fr/DSIR/DSIR.html) (74).

Top strand oligo: CTAGCAGTAGCTGGAGTACAACTTCAACATAGTTATATTCAAGCATATGTTGAAGTTGTACTCCAGCTGCG.

Bottom strand oligo: AATTCGCTGTTGAAGTTGTACTCCAGCTTATGCTTGAATATAACTAAGCTGGAGTACAACTTCAACAACTG.

After annealing the top and bottom strand oligoes, the product was inserted into the VALIUM22 vector, which had been previously linearized by NheI and EcoRI double restriction (New England Biolabs, lpswich, Massachusetts). The resulting plasmids were transformed, miniprepped with QIAprep Spin Miniprep Kit (QIAGEN, Hilder, Germany) and injected into fly strain *y sc v nanos-integrase; attP40* for stable transgene integration (75).

### UAS-Ndg^FL^.GFP

The coding sequence of Nidogen was amplified by PCR from 3rd instar larval cDNA with forward primer:

GGGGACAAGTTTGTACAAAAAAGCAGGCTTCATGCCGACCTTCGGCAGTAAGTTGC

and reverse primer:

GGGGACCACTTTGTACAAGAAAGCTGGGTC GTAGCCAGGCGCCAGCACGG.

The resulting product was cloned into PDONR211 (Life Technologies, Carlsbad, California) using Gateway BP Clonase Enzyme Mix (Life Technologies) to obtain PDONR211-Ndg, which was finally transferred into destination vector PTWG-1076 (UAST C-terminal GFP, *Drosophila* Carnegie Vector collection) plasmid by using Gateway LR Clonase Enzyme Mix (Life Technologies). Transgenic lines were obtained through standard P-element transgenesis (Spradling and Rubin, 1982).

### Other Nidogen constructs

For the structure/function analysis of Ndg domains, deletion of specific domains of Ndg was achieved by PCR-amplifying plasmid PDONR211-Ndg with the appropriate combinations of the following primers:

Ndg^G1^.GFP-Forward: TGAGAACGAGGACCCAGCTTTCTTGTACAAAG
Ndg^G1^.GFP-Reverse: AAGCTGGGTCCTCGTTCTCAATGGGAGCCAC
Ndg^G2^.GFP-Forward: GGTCAGCGGAGCTAATGATCAACCTATCCGAGTG
Ndg^G2^.GFP-Reverse: GATCATTAGCTCCGCTGACCAGGATCACCGAG
Ndg^G3^.GFP-Forward: CAGCGGACAGCGTCCCATTTCGGTGGCCC
Ndg^G3^.GFP-Reverse: AAATGGGACGCTGTCCGCTGACCAGGATCAC
Ndg^Rod^.GFP-Forward: GACATTACGTGACCCAGCTTTCTTGTAC
Ndg^Rod^.GFP-Reverse: AAGCTGGGTCGACATTACGTCCGTTTAG
Ndg^RodG3^.GFP-Forward: GGTCAGCGGAAACGATGGTACCGCCGATTG
Ndg^RodG3^.GFP-Reverse: TACCATCGTTTCCGCTGACCAGGATCACCGAG
Ndg^G1L^.GFP-Forward: GTCCTGCCTGTACGACCCAGCTTTCTTGTACAAAG
Ndg^G1L^.GFP-Reverse: CTGGGTCGTACAGGCAGGACTTTCCATTGCC
Ndg^G1LG2^.GFP-Forward: AAATGGGACGCAGGCAGGACTTTCCATTGCC
Ndg^G1LG2^.GFP-Reverse: CTGGGTCGTAATCGTTGCAGGCATTCGATTCGGG
Ndg^ΔG3^.GFP-Forward: GACATTACGTGACCCAGCTTTCTTGTAC
Ndg^ΔG3^.GFP-Reverse: AAGCTGGGTCGACATTACGTCCGTTTAG
Ndg^ΔG2R^.GFP-Forward: GTCCTGCCTGCGTCCCATTTCGGTGGCCCAG
Ndg^ΔG2R^.GFP-Reverse: AAATGGGACGCAGGCAGGACTTTCCATTGCC
Ndg^ΔG2^.GFP-Forward: GTCCTGCCTGAACGATGGTACCGCCGATTG
Ndg^ΔG2^.GFP-Reverse: TACCATCGTTCAGGCAGGACTTTCCATTGCC
Ndg^ΔG1^.GFP-Forward: GGTCAGCGGAGAGCAGAACGTGAGGTCTCCC
Ndg^ΔG1^.GFP-Reverse: CGTTCTGCTCTCCGCTGACCAGGATCACCGAG
Ndg^G2R^.GFP-Forward: GGTCAGCGGAGCTAATGATCAACCTATCCG
Ndg^G2R^.GFP-Reverse: GATCATTAGCTCCGCTGACCAGGATCACCG
Ndg^L^.GFP-Forward: GGTCAGCGGAGAGCAGAACGTGAGGTCTCC
Ndg^L^.GFP-Reverse: CGTTCTGCTCTCCGCTGACCAGGATCACCG
Ndg^ΔR^.GFP-Forward: GAATGCCTGCCGTCCCATTTCGGTGGCCCA
Ndg^ΔR^.GFP-Reverse: AAATGGGACGGCAGGCATTCGATTCGGGGG

The resulting PCR reactions were incubated with 10 units of DMT enzyme (TransGen Biotech, Beijing, China) at 37°C for 1 hour to digest the original templates. After digestion, PCR products were transformed into DMT competent cells (TransGen Biotech, Beijing, China). Colonies were validated by sequencing. Transgenic lines were obtained through standard P-element transgenesis (Spradling and Rubin, 1982).

### Generation of deficiencies removing the gene Ndg

The Mi{ET1}Ndg^MB12298^ transposon was used in a Blm mutant background to generate deficiencies by imprecise excision of the transposon. In these mutants, homologous recombination DNA reparing enzymes are compromised, thus increasing the events of non-homologous recombination DNA repair. Non-homologous recombination DNA repair increases the chances of generating DNA deficiencies (Witsell et al., 2009). We selected 132 Blm mutant males carrying the Mi{ET1}Ndg[MB12298] transposon and crossed them to w; Gla/CyO females. The offspring of this cross rendered a 110 EGFP negative males that were crossed to the Df(2R)BSC281 deficiency. 6 out of the 110 males did not complement the deficiency and were selected for further molecular characterization with the following primers from the Ndg genomic region. PCR primers were used as follows: (5’-3’)

Ndg primer1-Forward: GTGTGGACTCGGTGTGACTG
Ndg primer1-Reverse: ACTTCGAACAGCCAGACTCC
Ndg primer2-Forward: CCTTCGGCAGTAAGTTGCTC
Ndg primer2-Reverse: GTGCTGTTGGACAGACAACG
Ndg primer3-Forward: CGATCAAGCGGCGCAATATC
Ndg primer3-Reverse: CCAACATGCCACAATGGGTG
Ndg primer4-Forward: GTCTGAGTGGTTTCGGCAC
Ndg primer4-Reverse: TTTGCTTAAAGTGGGTGTTGC
Ndg primer5-Forward: CCATTGTGGCATGTTGGATA
Ndg primer5-Reverse: TGTTTCGAAGGCGATACTCA
Ndg primer6-Forward: AAACTGAAAAAGCGGGGAAT
Ndg primer6-Reverse: TTAATCAGTGCACCGCAGAG
Ndg primer7-Forward: GATGAAGGAGGCAAAGCAAG
Ndg primer7-Reverse: TTTTCATCTGCAGTGCGTTC
Ndg primer8-Forward: GAGGAGCAGATACCCCAACA
Ndg primer8-Reverse: CAGTGCCGTCATATTTGGTG
Ndg primer9-Forward: GGATTCAGAGGCGATGGATA
Ndg primer9-Reverse: GACCAGTTCCGTCCAGGTTA
Ndg primer10-Forward: TTTCTGCCAGTTTTCGCTTT
Ndg primer10-Reverse: CGTGTTGTTGGATTGTGGAG
Ndg primer11-Forward: GTGCTGTGCCTCAGATGAAA
Ndg primer11-Reverse: GGGAACCCAATGTGCTTAGA
Ndg primer12-Forward: TTACCTTCACGCACGATCAG
Ndg primer12-Reverse: GGCTGCGGCATTAGAGATAC

### Generation of *Ndg* mutants with CRISPR/Cas9

Four sgRNAs were designed for generating *Ndg* null mutant lines (76). sgRNAs and cas9 mRNA were injected into *w^1118^* embryos. *Ndg* deletions in the germ line *Ndg^1^* and *Ndg^2^* were selected by sequencing by Beijing Fungene Biotechnology (Beijing, China).

sgRNA1: GAGAGATACACAAGTCAGGAAGG
sgRNA2: CCAGCCCTTTCCGCTGGAATATGC
sgRNA3: GCGGCCTTCTACTCGAACGTGG
sgRNA4: GCCATTTGCAAGTGGGACTCGG

For assessment of Ndg mRNA expression in *Ndg^1^* mutants was assessed by quantitative real-time PCR. RNA was extracted using TRIzol reagent (Life technologies, USA). cDNA was synthesized from 2 μg of RNA with PrimeScriptTM RT-PCR Kit (Takara, Kyoto, Japan). Analysis was performed in a CFX96 Touch system (Bio-Rad, California, USA) using iTaq Universal SYBR Green Supermix (Bio-Rad). *rp49* was used as a reference for normalization. Three experiments per genotype were averaged. The following intron-spanning pairs of primers were used:

Ndg primerA-Forward: GAGCAGTACGAGCAGCT
Ndg primerA-Reverse: CGAGTAGAAGGCCGCTAT
Ndg primerB-Forward: ATCCATATCCTGAGGAGCAGAT
Ndg primerB-Reverse: GGTGCAGGTGTAGCCAT
Ndg primerC-Forward: AGTGCCGTTCGACCAATT
Ndg primerC-Reverse: GACAATCAGGAAGTCAGAGT
Ndg primerD-Forward: GACTCAGCAAAGGATACCAT
Ndg primerD-Reverse: CAGTCCGACCAGAACAGTT
rp49 primer-Forward: GGCCCAAGATCGTGAAGAAG
rp49 primer-Reverse: ATTTGTGCGACAGCTTAGCATATC

To generate Ndg mutants carrying a deletion of the rod and G3 domains, one single guide (sgRNA) target was designed in the 5^th^ exon of Ndg:

sgRNA5: GGGGAATGCCGATGCCCCTATGG

The sgRNAs were cloned in the PCFD3 vector as previously described in (77) and http://www.crisprflydesign.org/plasmids/. Transgenic gRNA flies were created by the Best Gene Company (Chino Hills, USA) using either *y sc v P{nos-phiC31\int.NLS}X; P{CaryP}attP2* (BDSC 25710) or *y v P{nos-phiC31\int.NLS}X; P{CaryP}attP40* (BDSC 25709). Transgenic lines were verified by sequencing by Biomedal Company (Armilla, Spain). Males carrying the sgRNA were crossed to females either act-Cas9 or nos-Cas9 and the progeny was screened for the v+ch- eye marker. To identify CRISPR/Cas9-induced mutations, genomic DNA was isolated from flies and sequenced using the following primers: (5’-3’)

Ndg primerg5-Forward: GCGAAGTTTGGGAGAACGGA
Ndg primerg5-Reverse: ACAGTATCTCACTCAGATCGGC

### Immunohystochemistry and imaging

For generation of anti-Nidogen antibody, rabbits were immunized with epitope CTYVQEFDGERNADLIPC by Bio-med Biotechnology (Beijing, China). Embryos, fat bodies, wing imaginal discs and ovaries were stained using standard procedures and mounted in DAPI-Vectashield (Vector Laboratories, Burlingame, California). The following primary antibodies were used: rabbit anti-Ndg (1:2000, this study), chicken anti-betagalactosidase (1:500, AbCam, Cambridge, UK), chicken anti-GFP (1:500, AbCam), rabbit anti-Ndg (1:100, (49). Secondary antibody is IgG conjugated to Alexa-555, IgG conjugated to Alexa-488 and Alexa 549 (1:200, Life technologies). For lipid droplet staining, L3 larvae were turned inside out and fixed in 4% PFA for 20 minutes, washed twice in PBS and then incubated in a 1:1000 dilution in PBS of 1 mg/ml BODIPY 493/503 stock (Life Technologies) for 30 minutes, followed by two 10-min washes in PBS and mounting in DAPI-Vectashield (Vector Laboratories). Confocal images were obtained using a Leica (Wetzlar, Germany) SP2 microscope or a Zeiss (Oberkochen, Germany) LSM780 microscope equipped with a Plan-Apochromat 63X oil objective (NA 1.4). Eggs and pupae were imaged in a Leica M125 stereoscope. All images were processed with Adobe Photoshop and ImageJ.

### Quantification

For quantification of egg laying (Fig. S4A), five 2-day old virgins were transferred to fresh vials daily for ten days and the eggs laid on each vial counted. Three such experiments were conducted per genotype.

For calculation of egg aspect ratio (Fig. S4B; Frydman and Spradling, 2001), length and width of eggs were measured on images using the line tool in FIJI-ImageJ. Aspect ratio is defined as egg length divided by width).

In chill comma recovery time assays (Fig. S4D; Gilbert et al., 2001), 2-day old females were placed into 10 mL tubes. These tubes were submerged into an ice-water bath for 2 hours, resulting in paralyzed flies. The amount of time required for a fly at room temperature to stand after becoming paralyzed in this way was measured.

For quantification of fluorescence intensity of different Ndg.GFP constructs in fat body BM (Fig. 6B), GFP signal was measured on 4-6 confocal images per genotype using FIJI-ImageJ. Each measurement represents mean value intensity inside a 500 μm^2^ square drawn on a flat portion of BM of an individual fat body cell, avoiding measuring intensity in cell contacts.

VNC length was measured on confocal images using the segmented line tool of FIJI-ImageJ.

### Statistical analysis

Graphpad Prism software was used for graphic representation and statistical analysis. For statistical comparisons of fluorescence intensity in Fig. 6B and 6C, unpaired Student’s t tests were used in *LanA^i^*+*Ndg^ΔG3^. GFP*, *Cg25C^i^+Ndg^FL^. GFP*, *Cg25C^i^+Ndg^ΔG1^. GFP* and *Cg25C^i^+Ndg^ΔG2^. GFP* experiments (data passed D’Agostino & Pearson normality tests and F-tests for equal variance). Student’s t tests with Welch’s correction were used for *LanA^i^+Ndg^FL^. GFP*, *LanA^i^+Ndg^ΔG1^. GFP* and *LanA^i^+Ndg^ΔG2^. GFP* experiments (data passed D’Agostino & Pearson normality tests, but not F-tests for equal variance). A non-parametric Mann-Whitney test was used in *Collagen IV^i^+Ndg^ΔG1^. GFP* experiment (data did not pass D’Agostino & Pewrson normality test). For comparisons of VNC length in Fig. 8A, we performed non-parametric Mann-Whitney tests. For egg production curves in Figure S4A, we conducted non-parametric Kolmogorov-Smirnov tests. For comparison of aspect ratio in Figure S4B, we performed unpaired two-tailed Student’s t tests. For comparison of chill comma recovery time in Figure S4C, Student’s t-tests with Welch’s correction were used. Significance of statistical tests is reported in graphs as follows: * * * * (p < 0.0001), * * * (p < 0.001), * * (p < 0.01), * (p < 0.05), n.s. (p > 0.05).

## Acknowledgments

We thank Bloomington Stock Center for fly stocks and for sharing valuable reagents and the Developmental Studies Hybridoma Bank from the University of Iowa (USA) and Dr A. Holz for antibodies.

## Supplemetal Figure Legends

**Fig S1. Fat body adipocytes and blood cells are the main source of BM components in the larva.**

Confocal images showing the localization of Collagen IV (Vkg.GFP, green), Perlecan (Trol.YFP, yellow) and Laminin (LanB1.sGFP, cyan) in different tissues of the third instar larva. Images compare control tissues (+) with tissues from larvae where expression of the corresponding fluorescence protein fusion has been knocked down through Cg-GAL4-driven iGFPi (*Cg> isGFPi*). Disappearance of the corresponding signal from BMs is observed, with the exceptions indicated by hollow arrows (partial reduction) and asterisks or filled arrows (no reduction). Nuclei stained with DAPI (white).

**Fig S2. Generation of deletions removing the *Ndg* gene.**

(A) Schematic representation of the deficiencies generated in the Ndg region by imprecise excision of the Mi{ET1}Ndg[MB12298] transposon (see Materials and Methods). (B) Colour-code diagram picture of the primer pairs used for molecular characterization of the deficiencies described in (A) (see Materials and Methods).

**Fig S3. Ndg accumulation in *Ndg*^ΔRod-G3.1^ homozygous mutant embryos.**

(A-C) Confocal images showing stage 16 embryos stained with two different anti-Ndg (red) antibodies. Images compare control embryos (A) with *Ndg*^ΔRod-G3.1^ embryos (B, C). (A-B) Stage 16 control (A) and *Ndg*^ΔRod-G3.1^ (B) mutant embryos stained with an anti-Ndg antibody (NDG-B) that recognizes the region between the second G2 domain up to the fourth EGF repeat of Ndg (Wolfstetter, 09). (B) While *Ndg*^ΔRod-G3.1^ embryos do not show any Ndg staining in embryonic BMs, expression in chordotonal organs is unchanged (arrowheads). (C) *Ndg*^ΔRod-G3.1^ mutant embryos stained with an anti-Ndg antibody that recognizes an epitope in the Rod domain (this work) do not show any Ndg expression.

**Fig S4. Elimination of Ndg affects egg deposition, adult flight muscle BM and chill coma recovery.**

(A) Quantification of eggs laid by wild type (*w^1118^*), *Ndg^1^* and *Ndg^2^* virgin females. The curves join mean values of three experiments (individual dots). Differences with the wild type are significant in Kolmogorov-Smirnov tests (**: p<0.01). (B) Images of eggs laid by wild type (*w^1118^*), *Ndg^1^* and *Ndg^2^* flies and graph quantifying egg aspect ratio (length/width). Each dot in the graph is a measurement from a single egg. Differences with the wild type were not significant in unpaired two-tailed Student’s t tests. (C) Images of the BM (Vkg.GFP in green) of adult flight muscles, showing the BM is broken in *Ndg^1^* mutants. (D) Quantification of chill coma recovery time in adult female control flies, *Ndg^1^* mutant, *Ndg^2^* mutant, *Ndg^1^/Df(2R)BSC281* and *Ndg^2^*/*Df(2R)BSC281*. Each dot in the graph is a measurement from a single fly. Differences with the wild type were significant in two-tailed t tests with Welch’s correction (****: p<0.0001). (B, D) Horizontal lines represent mean values.

**Fig S5. Regulation of Ndg.sGFP incorporation into BMs by laminins, Col IV and Perlecan.**

Confocal images of the fat body BM showing localization of Ndg (Ndg.sGFP, green), Laminin (LanB1.sGFP, cyan), Collagen IV (Vkg.GFP, green) and Perlecan (Trol.YFP). Images show fat body from wild type larvae (upper panels) and larvae where *LanA, LanB2* or *Cg25C* have been knocked down under control of Cg-GAL4.

